# Single-cell RNA sequencing identifies SLC6A3 as a biomarker and prognostic marker in clear cell renal cell carcinoma

**DOI:** 10.1101/2023.08.31.555693

**Authors:** Sathiya Pandi Narayanan, Ramani Gopal, Sebastian Arockia Jenifer, Tariq Ahmad Masoodi

## Abstract

**Background:** The SLC6A3 gene encodes dopamine protein and is a member of the sodium and chloride-dependent neurotransmitter transporter family. While the role of SLC6A3 in Parkinson’s disease is well established, its function in cancer, especially in clear cell renal cell carcinoma (ccRCC), remains unclear.

**Methods:** To investigate the expression and function of SLC6A3 in ccRCC, we conducted a study using single-cell transcriptomics and bulk RNA sequencing data. We analyzed advanced ccRCC single-cell RNA sequencing profiles, bulk RNA sequencing, and microarray data to assess the expression of SLC6A3 in tumor cells, benign kidney tubule cells, and immune cells.

**Results:** Our analysis showed that SLC6A3 expression is specific to ccRCC tumor cells and is not present in benign kidney tubule cells or immune cells of benign kidney and kidney tumors. Further, we found an elevated expression of SLC6A3 in ccRCC tumors compared to the benign kidney. Receiver operating characteristics analysis suggests that SLC6A3 is highly sensitive and specific to ccRCC. Additionally, we found a correlation between HNF4A signaling and SLC6A3 expression in two independent mRNA expression profiles. Interestingly, elevated expression of SLC6A3 is a predictor of better overall and progression-free survival of ccRCC patients.

**Conclusions:** Our findings suggest that SLC6A3 is a potential diagnostic and prognostic marker for ccRCC. The study highlights the importance of understanding the role of SLC6A3 in cancer and provides new insights into ccRCC diagnosis and treatment.

## Background

The kidney is a highly complex organ in the human body that performs various functions necessary for homeostasis (1–3). The kidney consists of a broad range of specialized cell types which are functionally diverse and spatially located in different anatomical segments (4, 5). Molecular alterations in the kidney cell types often lead to nephrological disorders and renal tumors (6, 7). Renal cancers are one of the significant causes of mortality. They are characterized by heterogeneous mRNA expression, DNA copy number, mutation, and epigenomic alterations, which may independently or together promote tumorigenesis (8, 9). In addition, the cellular composition of renal tumors and their immune microenvironment varies between renal cancer subtypes and patients (10, 11). So, understanding kidney tumor cell types and identifying the gene regulatory networks associated with cellular functions may provide deep molecular insights into renal cancers. Earlier, kidney tumor cell types have been annotated or defined by morphological appearance, spatial position, and a few selected marker genes expression (12, 13). Over the past two decades, the transcriptional landscape of benign and malignant kidney samples has been charted largely from bulk RNA sequencing, which quantitates the average expression of genes across a heterogeneous cell population (14, 15). However, recent single-cell transcriptomics (scRNA) technology quantitates mRNA in each cell and is based on the comprehensive gene expression pattern and refines the cellular identity. In addition, scRNA sequencing is also helpful in identifying novel cell types and marker genes in different cell types (16–18).

To date, many diagnostic and prognostic markers for various cancers, including clear cell RCC, are developed from bulk RNA sequencing data. However, there is a significant need for cell-type and tumor-type-specific markers (9). Single-cell RNA sequencing data allows us to quantitate the marker gene expression in individual cell types. This study used single-cell RNA sequencing data for renal cell cancer and identified a tumor cell type-specific prognostic marker for clear cell RCC.

SLC6A3 is a dopamine transporter gene mainly involved in the transport of dopamine into the cell (19, 20). However, the role of SLC6A3 in cancer has not been well established. Previous bulk RNA sequencing studies identified over-expression of SLC6A3 in renal cell carcinomas (21, 22). In addition, SLC6A3 was identified as a circulating biomarker in gastric cancer (23). However, the expression pattern of SLC6A3 in a single-cell level in ccRCC has not been explored. In this study, we characterized the expression pattern of SLC6A3 in renal cancer and normal kidney cell types at the single-cell level and validated it in bulk RNA sequencing and microarray profiles.

## Methods

### TCGA data analysis

In our research, we utilized the UALCAN database (24, 25) to study the expression of SLC6A3 in various pan-cancer samples. To perform our analysis, we utilized the TCGA dataset, a widely used dataset in cancer research. We created a box plot to visualize the distribution of SLC6A3 expression using log2(TPM+1) normalized data. Furthermore, we identified the top 25 differentially expressed genes between normal and clear cell RCC using TCGA data in the UALCAN database. This allowed us to compare the expression of SLC6A3 between normal and clear cell RCC, which can help in understanding the role of SLC6A3 in RCC development. To further investigate the expression of SLC6A3, we plotted a human anatomy graphical representation using the GEPIA2 tool (26). This visualization enabled us to observe the distribution of SLC6A3 expression across various tissues and organs. Additionally, we used the GEPIA2 tool to plot the RCC subtype (clear cell RCC, chromophobe RCC, and papillary RCC) specific expression of SLC6A3. We matched TCGA normal and GTEx data and set the following parameters (Log2FC cutoff – 1; p-value cutoff – 0.01). This analysis allowed us to identify the specific expression pattern of SLC6A3 in different RCC subtypes, which can help in understanding the role of SLC6A3 in RCC development and progression.

### Single-cell RNA sequencing data analysis

The Human Cell Atlas is an extensive repository that contains a vast collection of single-cell RNA sequencing datasets provided by numerous scientists. In our research, we utilized this valuable resource to explore the expression pattern of SLC6A3 in renal cell carcinoma. We examined the data from a study called “Tumor and immune reprogramming during immunotherapy in advanced renal cell carcinoma,” which analyzed over 34,000 (27) cells using the UMAP clustering method. The clusters were grouped based on their cell lineage, including Lymphoid, Myeloid, Normal Tissue, and Putative Tumor. By examining the expression of SLC6A3, CA9, and RHCG (14) in the UMAP clustering, we were able to identify the expression patterns of SLC6A3 and CA9 (28) in the Human Kidney Organoids Atlas dataset. Additionally, we analyzed the expression of SLC6A3, CA9, and RHCG in adult kidney single-cell data. Through these analyses, we gained a comprehensive understanding of the expression patterns of SLC6A3 in different types of kidney cells and identified potential biomarkers for renal cell carcinoma. By utilizing the Human Cell Atlas database, we were able to identify new insights into the molecular mechanisms underlying renal cell carcinoma and pave the way for developing more effective therapies for this devastating disease.

### Microarray data analysis

In this study, microarray mRNA datasets were used to analyze gene expression in clear cell renal cell carcinoma (ccRCC) tumors and adjacent benign kidney samples. The datasets were obtained from the Gene Expression Omnibus database (GEO) (29, 30), which is a public repository of functional genomics datasets. Specifically, two datasets were used: GSE40435 (31) and GSE53757 (32). GSE40435 contains 101 ccRCC tumors and 101 benign adjacent kidney samples, while GSE53757 has 72 ccRCC tumors and 72 benign adjacent kidney samples. The microarray platform-specific probes were mapped to gene symbols with appropriate annotation files downloaded from the vendor. This allowed the researchers to match the probes to specific genes and determine their expression levels. To analyze the data, the expression values of genes with multiple probes were averaged. The datasets were then normalized, and log2 transformation was performed. This helped to reduce the effects of technical variability in the data and allow for better comparison of expression levels across samples. Finally, differential expression analysis was performed using BRB array tools.

### Receiver operating characteristics (ROC) analysis

In this study we used two microarray datasets to perform ROC (Receiver Operating Characteristic) analysis. The first dataset, GSE40435, contained 101 ccRCC tumors and 101 benign adjacent kidney samples. The second dataset, GSE53757, contained 72 ccRCC tumors and 72 benign adjacent kidney samples. These datasets provided a large number of samples for analysis, which helped to increase the statistical power of the study. To perform ROC curve analysis, the microarray data was first normalized to account for technical variability in the data. Normalization ensures that the expression levels of genes are comparable across different samples. The normalized data was then used to plot the ROC curves. The ROC curves were generated using the MedCalc statistical software. This software is commonly used for statistical analysis in biomedical research. The DeLong et al. methodology was selected in the MedCalc tool to plot the curve. This methodology is a statistical method that is commonly used to compare the accuracy of two diagnostic tests.

### *Insilico* pathway scanning analysis

In this study, we used microarray datasets GSE40345 and GSE53757 for insilico pathway scanning analysis. This involved analyzing gene expression data to identify the pathways that were activated in clear cell renal cell carcinoma (ccRCC) tumors. To perform this analysis, we downloaded gene signatures (**Table. S1**) representative of corresponding pathways from the molecular signature database (MSigDB) (33, 34). These gene signatures provide information about the specific genes that are involved in different pathways. We then used R-Bioconductor to quantitate the pathway scores (**Script. S1**). This involved first Z-normalizing the expression data to account for technical variability in the data. Next, the signature-specific genes were additively combined to get pathway activation scores (35, 36). This allowed us to determine the overall activation status of different pathways in ccRCC tumors. To visualize the results of this analysis, we plotted the signature scores and SLC6A3 expression as a heatmap using the Morpheus tool. This tool is commonly used for visualizing large datasets and allows for easy interpretation of complex data. Overall, insilico pathway scanning analysis provided insights into the molecular pathways that are activated in ccRCC tumors. This information can be used to identify potential therapeutic targets for this disease and develop more targeted treatments for patients.

## Results

### Over-expression of SLC6A3 in clear cell renal cell carcinoma

In this study, we investigated the expression pattern of SLC6A3 in TCGA pan-cancer expression data. We compared the expression levels of SLC6A3 in clear cell renal cell carcinoma (ccRCC) and other kidney tumors with their normal adjacent tissues. Our analysis revealed that ccRCC had the highest expression of SLC6A3 among all tumor samples, as shown in (Fig. 1A). Additionally, the normal adjacent tissues showed under-expression of SLC6A3 compared to the KIRC tissues, as shown in (Fig. 1B). However, other kidney tumors in TCGA data, such as chromophobe RCC (KICH) and Papillary RCC (KIRP), did not over-express SLC6A3 compared to their normal adjacent counterparts. As depicted in (Fig. 1C), KIRC exhibited multiple folds of overexpression of SLC6A3 compared to KICH and KIRP. Furthermore, we identified the top 25 differentially expressed genes between ccRCC and normal kidney tissues. Interestingly, SLC6A3 was found to be the topmost differentially expressed gene in ccRCC compared to normal kidney tissue, as shown in (Fig. 1D). This discovery allows us to further explore the SLC6A3 gene in ccRCC and its potential role in the development and progression of this disease. Overall, our findings provide important insights into the molecular mechanisms underlying ccRCC and highlight the potential of SLC6A3 as a biomarker for this disease.

**Figure 1.**
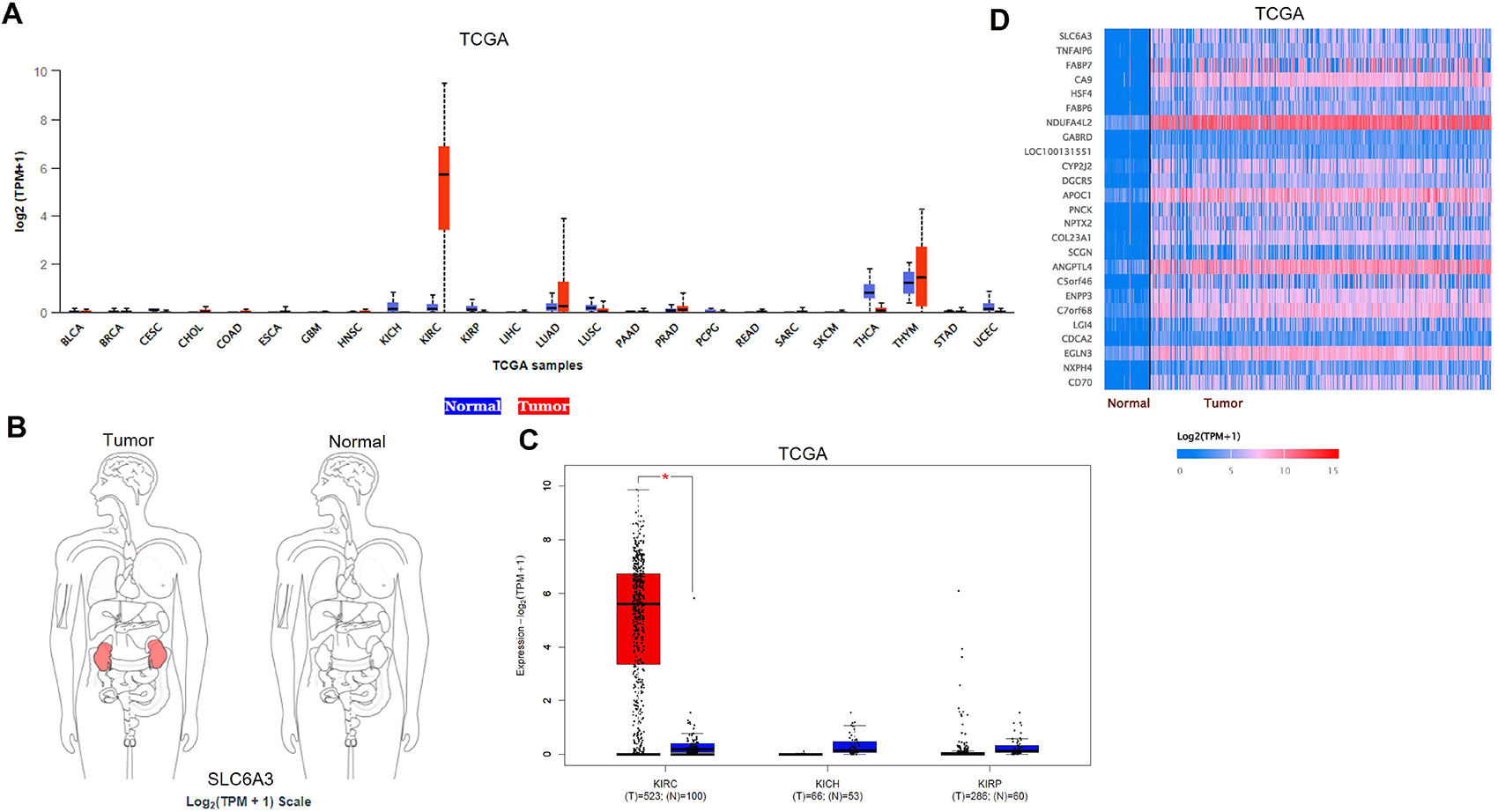
Expression pattern of SLC6A3 in pan-cancers. (A) Boxplot shows the expression pattern of SLC6A3 in different cancer types. (B) Schematic representation of SLC6A3 overexpression in tumor kidney compared to the normal kidney. (C) SLC6A3 expression pattern in major subtypes of renal cell carcinomas. (D) Heatmap shows the top 20 differentially expressed genes between clear cell renal cell carcinoma and normal kidney.

### The expression pattern of SLC6A3 in kidney cancer single-cell sequencing data

Next, we investigated the expression pattern of SLC6A3 in renal cancer single-cell sequencing data. We utilized the Human Cell Atlas data to query the SLC6A3 expression pattern and the UMAP plot to show the cell lineages present in advanced renal cell carcinoma. The UMAP plot classified four types of lineages, including lymphoid, myeloid, normal tissue, and putative tumor (**Fig. 2A**). To further investigate SLC6A3 expression, we checked the expression pattern of CA9, a well-known and widely used marker in ccRCC diagnosis. CA9 showed a high expression in the putative tumor cluster, indicating the presence of ccRCC tumor cells (**Fig. 2C**). Then we queried the SLC6A3 expression pattern and found that SLC6A3 was only expressed in the putative tumor cluster, and none of the other clusters showed SLC6A3 expression (**Fig. 2B**). Furthermore, we identified that RHCG, a marker for KICH, did not show expression in the putative tumor cluster, indicating that SLC6A3 is specifically expressed in ccRCC tumor cell types only (**Fig. 2D**). These results confirm that SLC6A3 is a promising diagnostic marker for ccRCC and can help in the development of targeted therapies for this disease.

**Figure 2.**
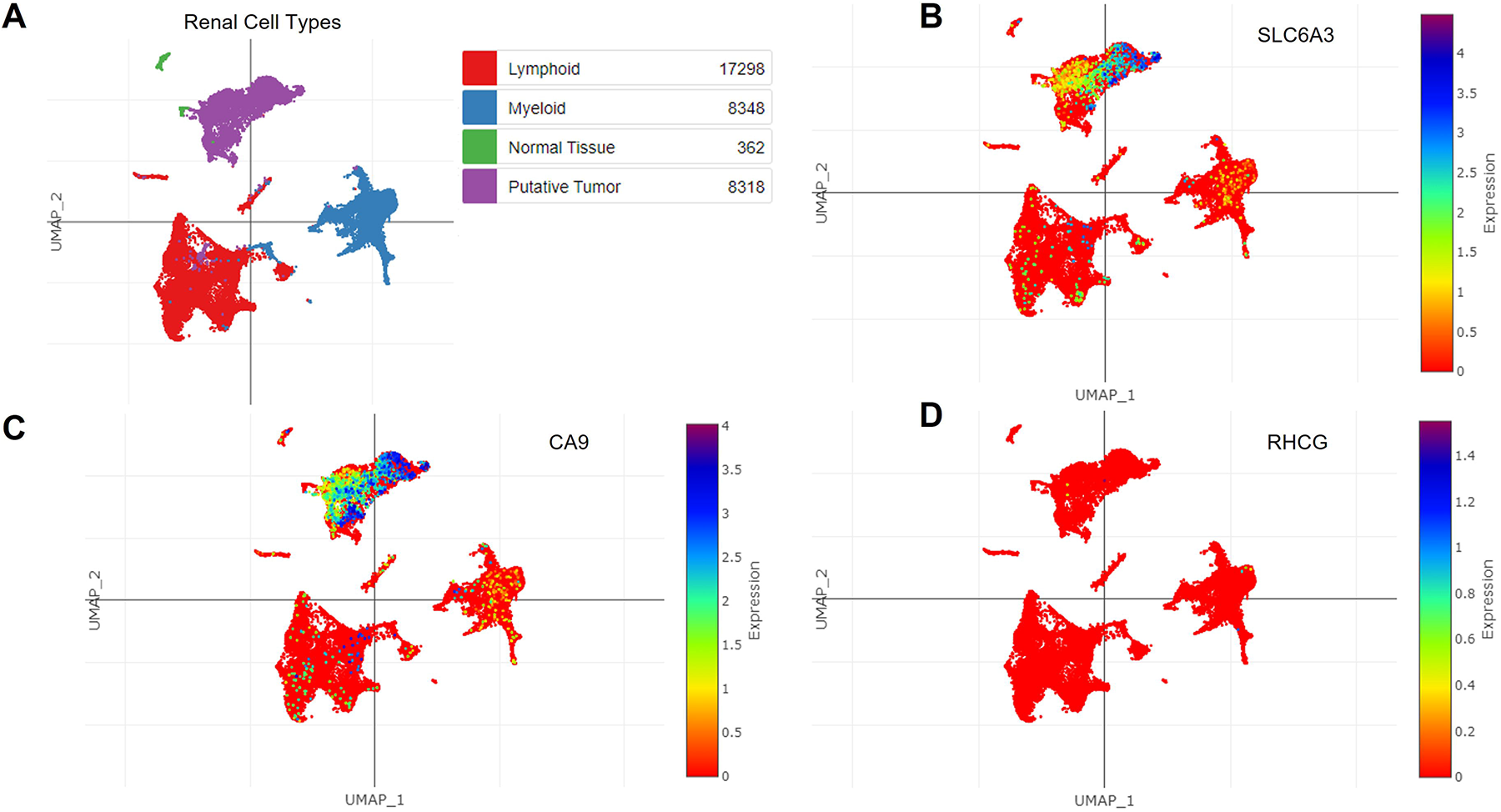
SLC6A3 expression in clear cell renal cell carcinoma single-cell RNA sequencing data. (A) The t-SNE plot shows the cell lineage types in the kidney single-cell RNA sequencing data. (B) The expression pattern of SLC6A3 is demonstrated in the t-SNE plot of kidney single-cell RNA sequencing data. (C and D). The expression pattern of CA9 and RHCG is shown in the t-SNE plot of kidney single-cell RNA sequencing data.

### The expression pattern of SLC6A3 in benign kidney Cell types

We investigated the expression of SLC6A3 in benign kidney single-cell atlas data and analyzed the UMAP data to identify multiple cell types present in the kidney, including epithelial lineage, distal nephron, collecting duct-like epithelial cells, glomerular cells, muscle-like cells, and endothelial cells. The UMAP plot visualized the distribution of these cells in the kidney cortex and medulla regions (**Fig. 3A**). We found that none of the cell types in the benign kidney cell atlas expressed CA9 and SLC6A3 (**Fig. 3B, C**). These results further confirm that the expression pattern of SLC6A3 is specific to tumor cells in ccRCC and is not expressed in any normal or immune cells of the kidney. This finding highlights the potential of SLC6A3 as a specific diagnostic marker for ccRCC and supports its use in developing targeted therapies for this disease. In conclusion, our investigation of SLC6A3 expression in benign kidney single-cell atlas data adds to the growing body of evidence supporting the specificity of this marker for ccRCC. The data suggests that SLC6A3 expression is limited to ccRCC tumor cells and is not expressed in any of the normal or immune cells of the kidney, providing further evidence for its potential as a specific diagnostic marker and therapeutic target for ccRCC.

**Figure 3.**
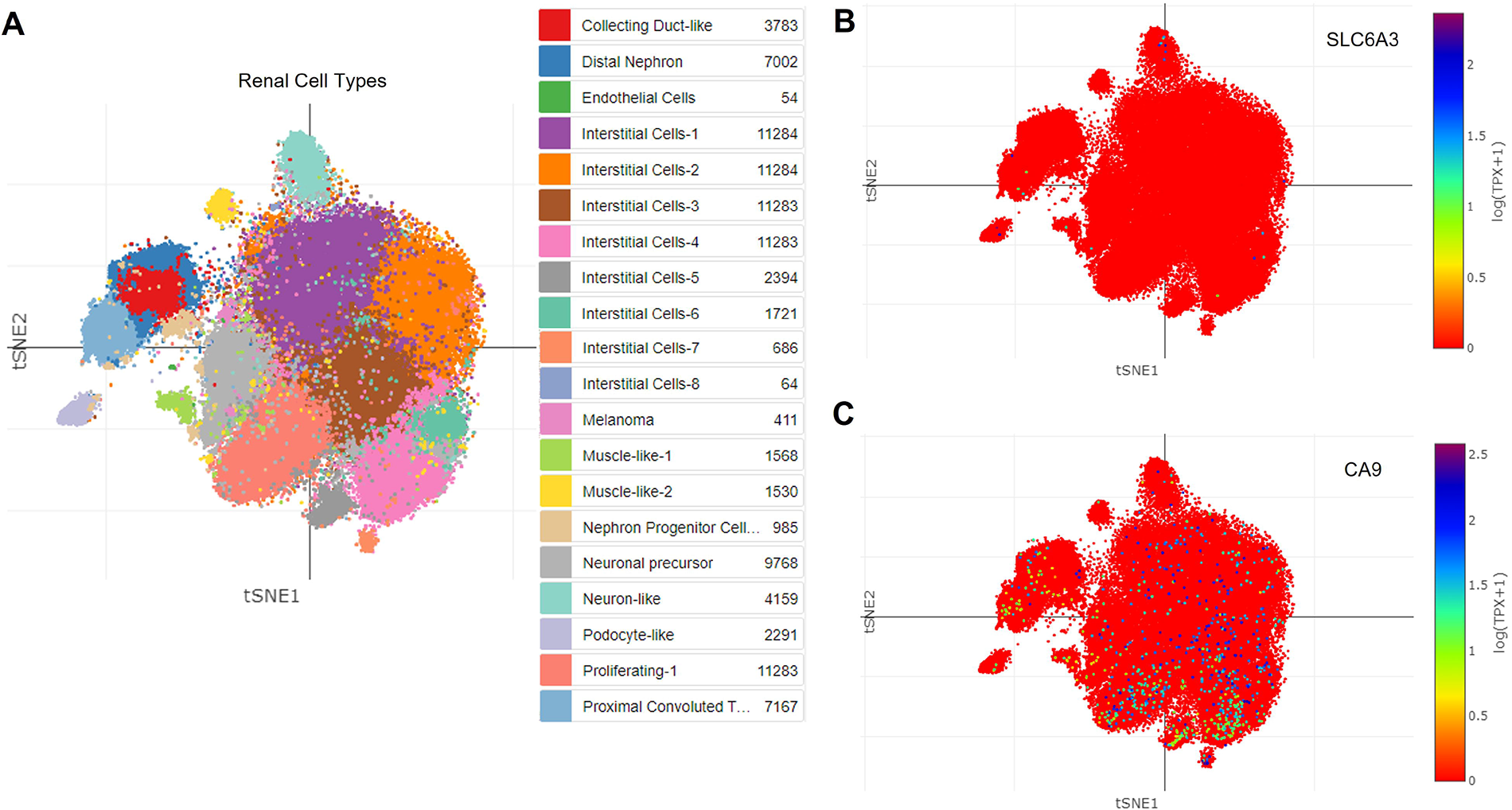
Expression pattern of SLC6A3 in benign kidney single-cell RNA sequencing data. (A) The t-SNE plot shows the various epithelial and immune cell types in the benign kidney single-cell RNA sequencing data. (B) The expression pattern of SLC6A3 is demonstrated in the t-SNE plot of benign kidney single-cell RNA sequencing data. (C). The expression pattern of CA9 is shown in the t-SNE plot of benign kidney single-cell RNA sequencing data.

### Investigation of SLC6A3 expression in ccRCC tumors

The investigation of SLC6A3 expression in multiple ccRCC expression datasets provided consistent results across all three independent datasets (**Fig. 4A, B**). The overexpression of SLC6A3 in ccRCC compared to normal kidney tissues was confirmed in all three datasets. Additionally, we performed stage-specific analysis of SLC6A3 expression in TCGA ccRCC expression data and found that SLC6A3 expression in stage IV is significantly lower than in stages I and II (**Fig. 4C**). To assess the diagnostic potential of SLC6A3, we conducted receiver operating characteristics analysis using the datasets GSE40435 and GSE53757. The analysis showed that SLC6A3 expression had high sensitivity and specificity for ccRCC tumors. In GSE40435, SLC6A3 expression showed 87.1% sensitivity and 97.0% specificity, while in GSE53757, it showed 93.1% sensitivity and 95.8% specificity (**Fig. 4D**). These findings indicate that SLC6A3 is a highly sensitive and specific marker for ccRCC tumors and has the potential to be used in diagnostic and therapeutic applications. Overall, our analysis of SLC6A3 expression in multiple ccRCC datasets provides strong evidence for its specificity and potential use as a diagnostic marker for this disease. The stage-specific analysis further supports the importance of SLC6A3 in the early stages of ccRCC. These findings suggest that further research is warranted to explore the therapeutic potential of SLC6A3 in the treatment of ccRCC.

**Figure 4.**
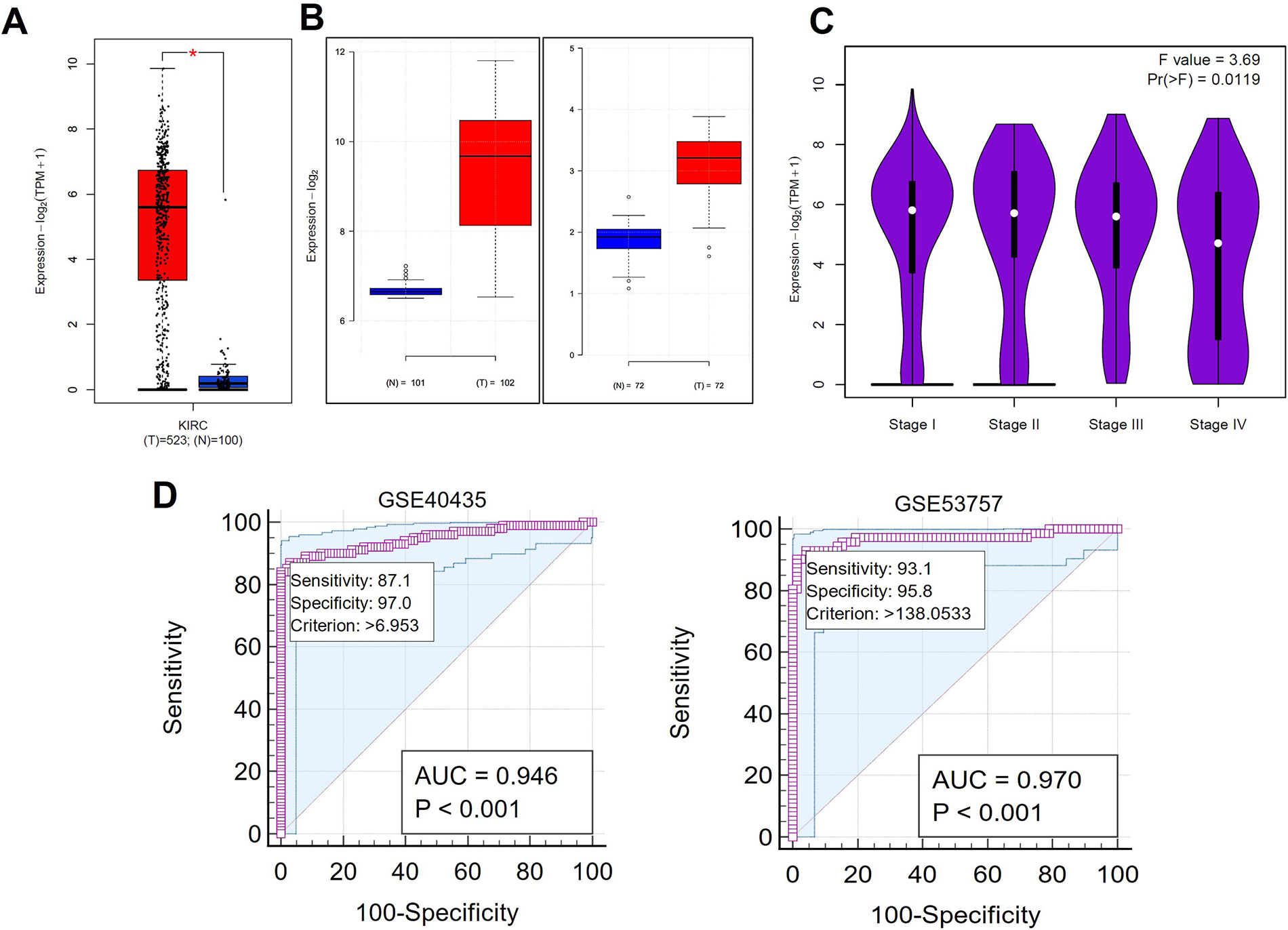
SLC6A3 expression pattern in various clear cell renal cell carcinoma mRNA expression datasets. (A and B) Boxplot shows the overexpression of SLC6A3 in clear cell renal cell carcinoma compared to the benign kidney. (C) Stage-specific expression of SLC6A3 in clear cell renal cell carcinoma. (D) Sensitivity and specificity analysis of SLC6A3 in predicting the clear cell renal cell carcinoma patients.

### SLC6A3 expression and signaling pathway correlation in ccRCC tumors

We conducted a gene signature-based pathway enrichment analysis in two different mRNA expression profiles of ccRCC tumor cohorts to identify the signaling pathway events correlated with SLC6A3 gene expression. We downloaded 32 different gene signatures from the molecular signature database and analyzed only tumor samples from the ccRCC cohort. The analysis provided normalized pathway activation scores for each gene set, which we plotted as a heatmap using the Morpheus tool. Our analysis showed that SLC6A3 expression is correlated with the HNF4A targets signaling pathway, which was strongly associated with SLC6A3 expression in GSE40345 and GSE53757 ccRCC mRNA expression profiles (**Fig. 5A, B**). Although our analysis provides a preliminary indication of the correlation between SLC6A3 expression and the HNF4A signaling pathway, further in vitro experimentation and validation in ccRCC tumor tissues are necessary to confirm the findings. These studies could provide a deeper understanding of the mechanisms underlying the involvement of SLC6A3 in the development and progression of ccRCC and may identify novel therapeutic targets for this disease.

**Figure 5.**
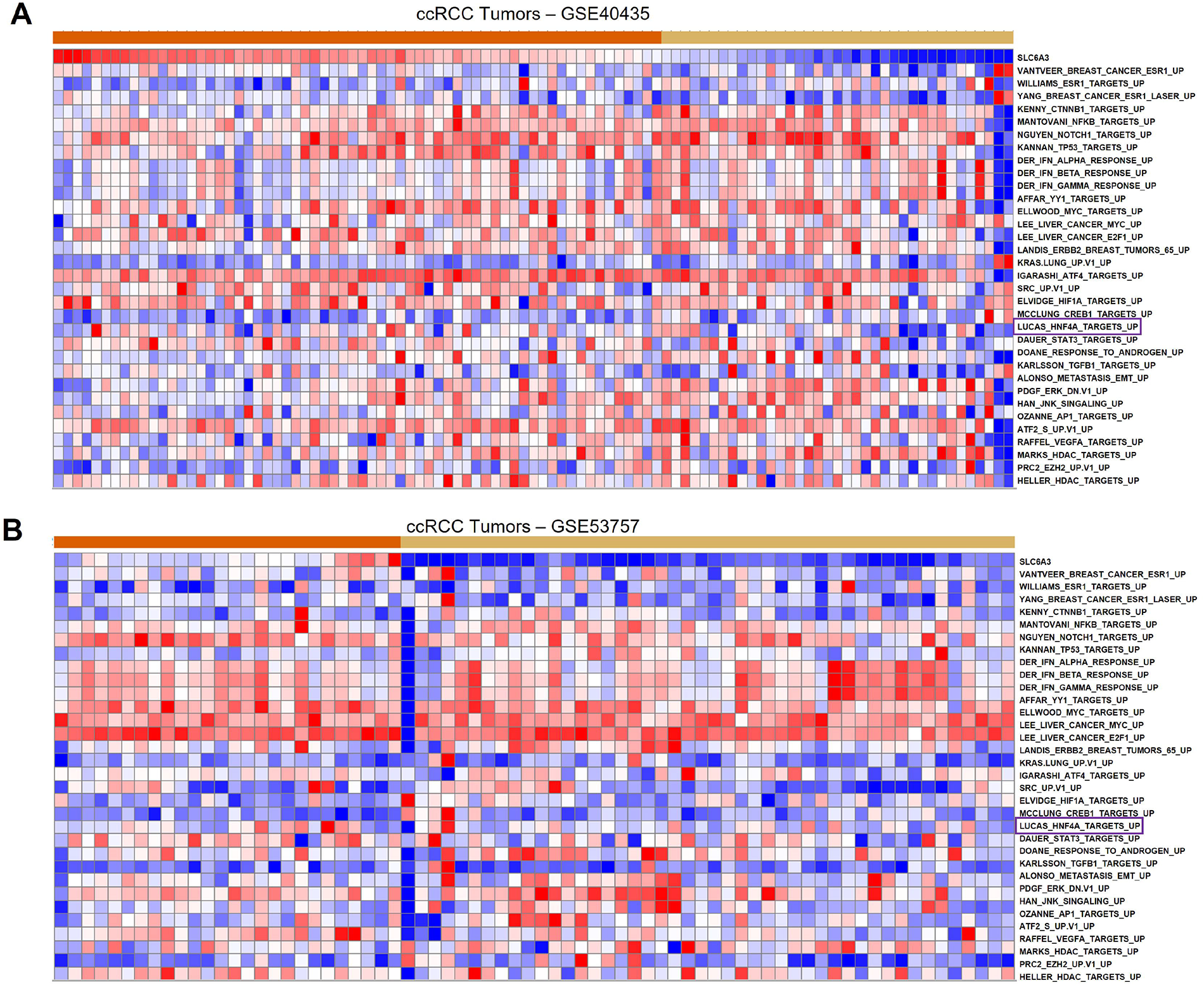
Identifying signaling pathways associated with SLC6A3 expression in clear cell renal cell carcinoma. (A) Heatmap shows the correlation of various signaling pathways associated with the SLC6A3 expression in clear cell renal cell carcinoma.

### Prognostic association of SLC6A3 in ccRCC tumors

Next, we investigated the prognostic significance of SLC6A3 in ccRCC using the TCGA ccRCC cohort. Our analysis revealed that higher expression of SLC6A3 was associated with better overall survival, while lower expression was associated with poorer overall survival. Furthermore, higher SLC6A3 expression was associated with better progression-free survival, while lower expression was associated with the worst progression-free survival in ccRCC patients (Fig. 6A, B). These findings indicate that SLC6A3 expression is not only a reliable diagnostic marker but also a potential prognostic indicator for ccRCC patients. The high sensitivity and specificity of SLC6A3 expression in predicting the survival outcomes of ccRCC patients can aid in the development of targeted therapies for the management of this disease.

**Figure 6.**
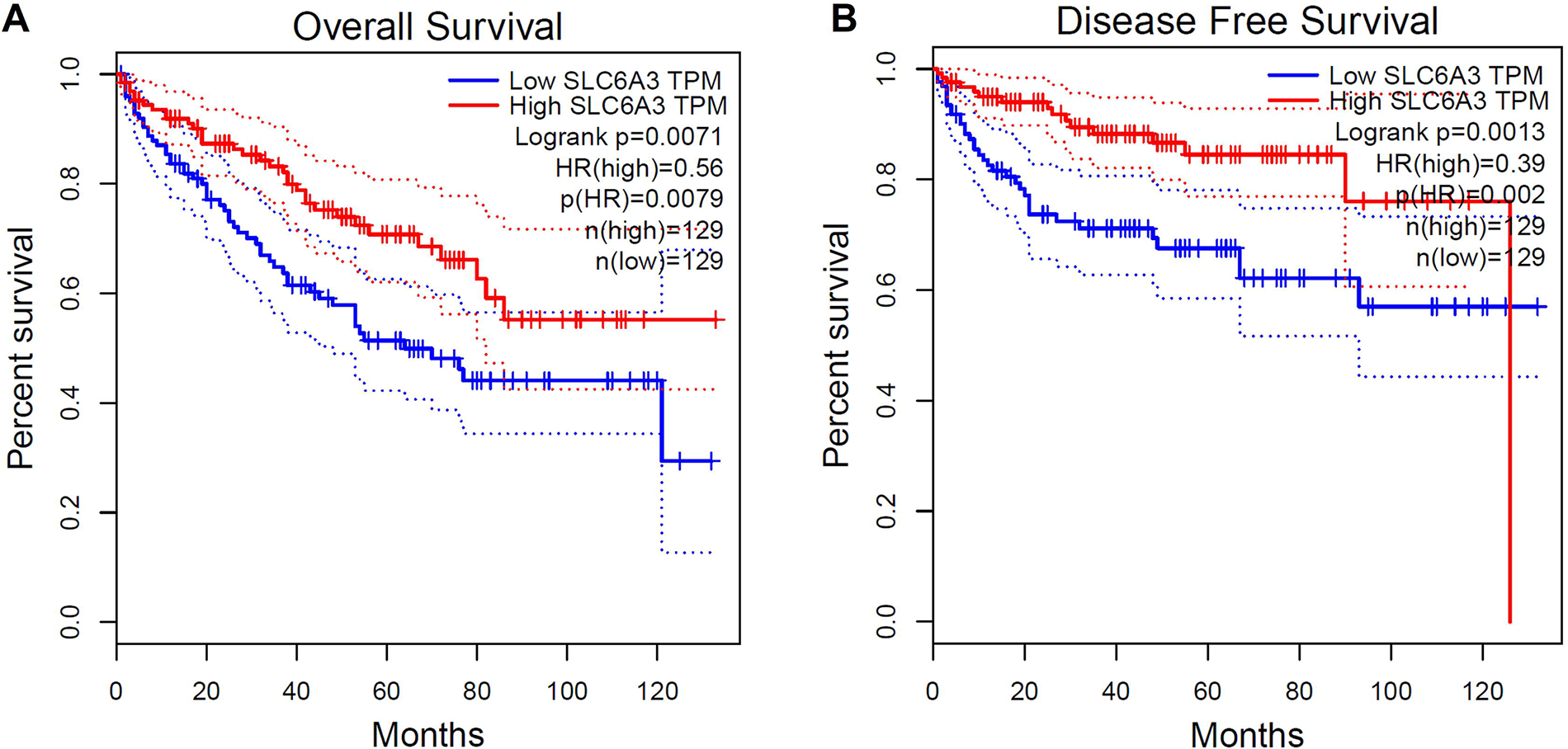
Kaplan-Meier survival analysis. (A and B) Kaplan-Meier survival curve analysis shows SLC6A3 expression associated with the survival of clear cell renal cell carcinoma patients.

## Discussion

Complementary experimental methodologies to investigate changes that underlie disease processes using either bulk tissue or single cells currently exist at transcriptomic, genomic, and epigenomic levels (37, 38). This transformative approach is now being used to create cell atlases of benign and diseased organs and tissues from humans, murine, and several other model systems. So much data has been generated and analyzed to identify novel cell type-specific, lineage-specific, and cancer-specific markers. However, the existing diagnostic and prognostic markers available for various cancer types have been extracted from the bulk RNA sequencing data (39, 40). However, the drawback of bulk RNA sequencing data is that the resultant expression data is from multiple cell types in the tissues. So, there is a strong need to identify novel markers from single-cell data for better diagnosis and therapeutic targeting in various cancers. In this study, we have utilized single-cell transcriptomics data to determine the sensitive and specific marker for the ccRCC tumors. First, we have identified the top 25 differentially expressed genes in ccRCC. Interestingly, we found that SLC6A3 is the top most gene on the list. In addition, pan-cancer analysis shows that SLC6A3 expression is precise to ccRCC compared to other tumor types. However, lung tumors and thymoma tumors show some expression, but they cannot be compared with SLC6A3 expression in ccRCC tumors. To date, two independent studies have reported the elevated expression of SLC6A3 in ccRCC tumors. Hansson et al. found that HIF-2α influences the expression of SLC6A3 in ccRCC cells, and they also said that hypoxia could induce SLC6A3 in normal renal cells (21). The other study by Schrödter et al. found that SLC6A3 mRNA is highly expressed in ccRCC tumors compared to normal kidney cells. Surprisingly, they also identified that the protein levels are higher in the normal tissue’s proximal tubule region than in tumor tissue (22). However, these two studies used bulk RNA expression data to identify the expression of SLC6A3 in ccRCC tumors. In this study, we used single-cell transcriptomics data to validate the expression pattern of SLC6A3 in ccRCC tumors. Remarkably, we found that SLC6A3 is only expressed in ccRCC tumor cells. None of the immune cells in the ccRCC shows SLC6A3 expression. In addition, none of the benign kidney cell types shows the expression of SLC6A3. Further, we wanted to evaluate the sensitivity and specificity of the SLC6A3 gene expression in ccRCC tumors. Receiver operating characteristics analysis shows that SLC6A3 is a precise marker for ccRCC with very extreme sensitivity and specificity. Next, we performed silico pathway scanning analysis using microarray datasets of ccRCC tumors. We found that excellent correlation between HNF4 signaling and SLC6A3 expression in two independent datasets. However, we did not find any association between HIF-1α signaling and SLC6A3 expression. Hansson et al. show that HIF-1α signaling is associated with the expression of SLC6A3 in ccRCC tumors. However, in silico method is a very subjective method. So, this identification must be validated and confirmed with in vitro experiments. In addition, we found an association between SLC6A3 expression and prognosis in ccRCC tumors. Schrödter et al. previously reported high protein expression of SLC6A3 in tumor tissues is associated with a shorter period of recurrence-free survival (22). In contrast, our data found that the higher expression of SLC6A3 is associated with better overall and progression-free survival. The discrepancy is due to the difference between the data used to quantitate the progression-free survival. Schrödter et al. used protein data to plot the survival graph, and this study used RNA expression to plot the survival graph (22). However, this needs to be revalidated in multiple cohorts for better confirmation. Together our results suggest that SLC6A6 is an independent and specific diagnostic and prognostic factor in ccRCC. SLC6A3 might become a valuable target for therapeutic options. Assessment of SLC6A3 expression in oral cancer samples might provide important information about patient’s clinical outcomes and therapeutic development.

## Conclusions

In conclusion, the use of single-cell transcriptomics data has enabled the identification of a sensitive and specific marker for clear cell renal cell carcinoma (ccRCC) tumors. The top most differentially expressed gene in ccRCC was found to be SLC6A3, which was shown to be expressed only in ccRCC tumor cells and not in immune cells or benign kidney cells. SLC6A3 was found to be a precise marker for ccRCC with high sensitivity and specificity, making it a valuable diagnostic and prognostic factor in ccRCC. The association between SLC6A3 expression and prognosis in ccRCC tumors was also studied, with conflicting results reported by different studies. The use of SLC6A3 as a therapeutic target in ccRCC warrants further investigation. Overall, the study highlights the importance of utilizing single-cell transcriptomics data to identify novel markers for better diagnosis and therapeutic targeting in various cancers.

## List of abbreviations

RCC: Renal Cell Carcinoma
ccRCC: Clear Cell Renal Cell Carcinoma
scRNA: Single Cell RNA sequencing
UALCAN: The University of ALabama at Birmingham CANcer data analysis Portal
TCGA: The Cancer Genome Atlas
GTEx: The Genotype-Tissue Expression
TPM: Transcript per Million
GEPIA2: Gene Expression Profiling Interactive Analysis
UMAP: Uniform Manifold Approximation and Projection

## Declarartion

### Ethics approval and consent to participate

Not Applicable

### Consent for publication

Not Applicable

### Availability of data and material

The single cell RNA sequencing datasets analysed during the current study are available in the Human Cell Atlas data portal repository. https://singlecell.broadinstitute.org/single_cell/study/SCP1288/tumor-and-immune-reprogramming-during-immunotherapy-in-advanced-renal-cell-carcinoma#study-summary.

The microarray datasets analysed during the current study are available in the GEO data repository.

https://www.ncbi.nlm.nih.gov/geo/query/acc.cgi?acc=GSE40435

https://www.ncbi.nlm.nih.gov/geo/query/acc.cgi?acc=GSE53757

The RNA sequencing datasets analysed during the current study are available in the UALCAN and GEPIA2 data repositories.

https://ualcan.path.uab.edu/ http://gepia2.cancer-pku.cn/

### Competing interests

The authors declare that they have no competing interests

### Funding

None to disclose.

### Authors’ contributions

SPN and TAM conceived the idea. SS and RG executed the data analysis. SAJ performed the statistical analysis and SS, RG, TAM and SPN written the manuscript.

## Acknowledgements

Not applicable

## Notes

### Competing Interest Statement

The authors have declared no competing interest.

## References

1. Robson L. The kidney--an organ of critical importance in physiology. J Physiol. 2014;592(18):3953–4.

2. Kariyanna SS, Light RP, Agarwal R. A longitudinal study of kidney structure and function in adults. Nephrol Dial Transplant. 2010;25(4):1120–6.

3. Solak Y. A longitudinal study of kidney structure and function in adults. Nephrol Dial Transplant. 2010;25(10):3457; author reply -8.

4. Balzer MS, Rohacs T, Susztak K. How Many Cell Types Are in the Kidney and What Do They Do? Annu Rev Physiol. 2022;84:507–31.

5. Woloshuk A, Khochare S, Almulhim AF, McNutt AT, Dean D, Barwinska D, et al. In Situ Classification of Cell Types in Human Kidney Tissue Using 3D Nuclear Staining. Cytometry A. 2021;99(7):707–21.

6. Alaghehbandan R, Perez Montiel D, Luis AS, Hes O. Molecular Genetics of Renal Cell Tumors: A Practical Diagnostic Approach. Cancers (Basel). 2019;12(1).

7. Testa U, Pelosi E, Castelli G. Genetic Alterations in Renal Cancers: Identification of The Mechanisms Underlying Cancer Initiation and Progression and of Therapeutic Targets. Medicines (Basel). 2020;7(8).

8. Ricketts CJ, De Cubas AA, Fan H, Smith CC, Lang M, Reznik E, et al. The Cancer Genome Atlas Comprehensive Molecular Characterization of Renal Cell Carcinoma. Cell Rep. 2018;23(1):313–26 e5.

9. Motzer RJ, Martini JF, Mu XJ, Staehler M, George DJ, Valota O, et al. Molecular characterization of renal cell carcinoma tumors from a phase III anti-angiogenic adjuvant therapy trial. Nat Commun. 2022;13(1):5959.

10. Zhao E, Li L, Zhang W, Wang W, Chan Y, You B, et al. Comprehensive characterization of immune- and inflammation-associated biomarkers based on multi-omics integration in kidney renal clear cell carcinoma. J Transl Med. 2019;17(1):177.

11. Diaz-Montero CM, Rini BI, Finke JH. The immunology of renal cell carcinoma. Nat Rev Nephrol. 2020;16(12):721–35.

12. Lindgren D, Eriksson P, Krawczyk K, Nilsson H, Hansson J, Veerla S, et al. Cell-Type-Specific Gene Programs of the Normal Human Nephron Define Kidney Cancer Subtypes. Cell Rep. 2017;20(6):1476–89.

13. Bachmann S, Kriz W, Kuhn C, Franke WW. Differentiation of cell types in the mammalian kidney by immunofluorescence microscopy using antibodies to intermediate filament proteins and desmoplakins. Histochemistry. 1983;77(3):365–94.

14. Skala SL, Wang X, Zhang Y, Mannan R, Wang L, Narayanan SP, et al. Next-generation RNA Sequencing-based Biomarker Characterization of Chromophobe Renal Cell Carcinoma and Related Oncocytic Neoplasms. Eur Urol. 2020;78(1):63–74.

15. Wach S, Taubert H, Weigelt K, Hase N, Kohn M, Misiak D, et al. RNA Sequencing of Collecting Duct Renal Cell Carcinoma Suggests an Interaction between miRNA and Target Genes and a Predominance of Deregulated Solute Carrier Genes. Cancers (Basel). 2019;12(1).

16. Zhang Y, Narayanan SP, Mannan R, Raskind G, Wang X, Vats P, et al. Single-cell analyses of renal cell cancers reveal insights into tumor microenvironment, cell of origin, and therapy response. Proc Natl Acad Sci U S A. 2021;118(24).

17. Liao J, Yu Z, Chen Y, Bao M, Zou C, Zhang H, et al. Single-cell RNA sequencing of human kidney. Sci Data. 2020;7(1):4.

18. Young MD, Mitchell TJ, Custers L, Margaritis T, Morales-Rodriguez F, Kwakwa K, et al. Single cell derived mRNA signals across human kidney tumors. Nat Commun. 2021;12(1):3896.

19. Reith MEA, Kortagere S, Wiers CE, Sun H, Kurian MA, Galli A, et al. The dopamine transporter gene SLC6A3: multidisease risks. Mol Psychiatry. 2022;27(2):1031–46.

20. Zhai D, Li S, Zhao Y, Lin Z. SLC6A3 is a risk factor for Parkinson’s disease: a meta-analysis of sixteen years’ studies. Neurosci Lett. 2014;564:99–104.

21. Hansson J, Lindgren D, Nilsson H, Johansson E, Johansson M, Gustavsson L, et al. Overexpression of Functional SLC6A3 in Clear Cell Renal Cell Carcinoma. Clin Cancer Res. 2017;23(8):2105–15.

22. Schrodter S, Braun M, Syring I, Klumper N, Deng M, Schmidt D, et al. Identification of the dopamine transporter SLC6A3 as a biomarker for patients with renal cell carcinoma. Mol Cancer. 2016;15:10.

23. Liu S, Cui M, Zang J, Wang J, Shi X, Qian F, et al. SLC6A3 as a potential circulating biomarker for gastric cancer detection and progression monitoring. Pathol Res Pract. 2021;221:153446.

24. Chen F, Chandrashekar DS, Scheurer ME, Varambally S, Creighton CJ. Global molecular alterations involving recurrence or progression of pediatric brain tumors. Neoplasia. 2022;24(1):22–33.

25. Zhang Y, Chen F, Chandrashekar DS, Varambally S, Creighton CJ. Proteogenomic characterization of 2002 human cancers reveals pan-cancer molecular subtypes and associated pathways. Nat Commun. 2022;13(1):2669.

26. Tang Z, Kang B, Li C, Chen T, Zhang Z. GEPIA2: an enhanced web server for large-scale expression profiling and interactive analysis. Nucleic Acids Res. 2019;47(W1):W556–W60.

27. Bi K, He MX, Bakouny Z, Kanodia A, Napolitano S, Wu J, et al. Tumor and immune reprogramming during immunotherapy in advanced renal cell carcinoma. Cancer Cell. 2021;39(5):649–61 e5.

28. Baniak N, Flood TA, Buchanan M, Dal Cin P, Hirsch MS. Carbonic anhydrase IX (CA9) expression in multiple renal epithelial tumour subtypes. Histopathology. 2020;77(4):659–66.

29. Barrett T, Wilhite SE, Ledoux P, Evangelista C, Kim IF, Tomashevsky M, et al. NCBI GEO: archive for functional genomics data sets--update. Nucleic Acids Res. 2013;41(Database issue):D991–5.

30. Edgar R, Domrachev M, Lash AE. Gene Expression Omnibus: NCBI gene expression and hybridization array data repository. Nucleic Acids Res. 2002;30(1):207–10.

31. Wozniak MB, Le Calvez-Kelm F, Abedi-Ardekani B, Byrnes G, Durand G, Carreira C, et al. Integrative genome-wide gene expression profiling of clear cell renal cell carcinoma in Czech Republic and in the United States. PLoS One. 2013;8(3):e57886.

32. von Roemeling CA, Radisky DC, Marlow LA, Cooper SJ, Grebe SK, Anastasiadis PZ, et al. Neuronal pentraxin 2 supports clear cell renal cell carcinoma by activating the AMPA-selective glutamate receptor-4. Cancer Res. 2014;74(17):4796–810.

33. Liberzon A, Birger C, Thorvaldsdottir H, Ghandi M, Mesirov JP, Tamayo P. The Molecular Signatures Database (MSigDB) hallmark gene set collection. Cell Syst. 2015;1(6):417–25.

34. Liberzon A, Subramanian A, Pinchback R, Thorvaldsdottir H, Tamayo P, Mesirov JP. Molecular signatures database (MSigDB) 3.0. Bioinformatics. 2011;27(12):1739–40.

35. Narayanan SP, Singh S, Gupta A, Yadav S, Singh SR, Shukla S. Integrated genomic analyses identify KDM1A’s role in cell proliferation via modulating E2F signaling activity and associate with poor clinical outcome in oral cancer. Cancer Lett. 2015;367(2):162–72.

36. Pandi NS, Suganya S, Rajendran S. In silico analysis of stomach lineage specific gene set expression pattern in gastric cancer. Biochem Biophys Res Commun. 2013;439(4):539–46.

37. Zhang Y, Wang D, Peng M, Tang L, Ouyang J, Xiong F, et al. Single-cell RNA sequencing in cancer research. J Exp Clin Cancer Res. 2021;40(1):81.

38. Jia Q, Chu H, Jin Z, Long H, Zhu B. High-throughput single-small es, Cyrillicell sequencing in cancer research. Signal Transduct Target Ther. 2022;7(1):145.

39. Joanito I, Wirapati P, Zhao N, Nawaz Z, Yeo G, Lee F, et al. Single-cell and bulk transcriptome sequencing identifies two epithelial tumor cell states and refines the consensus molecular classification of colorectal cancer. Nat Genet. 2022;54(7):963–75.

40. Zhang Z, Wang ZX, Chen YX, Wu HX, Yin L, Zhao Q, et al. Integrated analysis of single-cell and bulk RNA sequencing data reveals a pan-cancer stemness signature predicting immunotherapy response. Genome Med. 2022;14(1):45.

