## Supplemental File for "Single-cell RNA sequencing identifies SLC6A3 as a biomarker and prognostic marker in clear cell renal cell carcinoma"

**Table. S1.** List of gene signatures/gene sets used in this study. Gene sets were extracted from mSigDB. Totally 33 gene signatures utilized in this study.

| S. No. | Gene Signature |
|---|---|
| 1 | VANTVEER_BREAST_CANCER_ESR1_UP |
| 2 | WILLIAMS_ESR1_TARGETS_UP |
| 3 | YANG_BREAST_CANCER_ESR1_LASER_UP |
| 4 | KENNY_CTNNB1_TARGETS_UP |
| 5 | MANTOVANI_NFKB_TARGETS_UP |
| 6 | NGUYEN_NOTCH1_TARGETS_UP |
| 7 | KANNAN_TP53_TARGETS_UP |
| 8 | DER_IFN_ALPHA_RESPONSE_UP |
| 9 | DER_IFN_BETA_RESPONSE_UP |
| 10 | DER_IFN_GAMMA_RESPONSE_UP |
| 11 | AFFAR_YY1_TARGETS_UP |
| 12 | ELLWOOD_MYC_TARGETS_UP |
| 13 | LEE_LIVER_CANCER_MYC_UP |
| 14 | LEE_LIVER_CANCER_E2F1_UP |
| 15 | LANDIS_ERBB2_BREAST_TUMORS_65_UP |
| 16 | KRAS.LUNG_UP.V1_UP |
| 17 | IGARASHI_ATF4_TARGETS_UP |
| 18 | SRC_UP.V1_UP |
| 19 | ELVIDGE_HIF1A_TARGETS_UP |
| 20 | MCCLUNG_CREB1_TARGETS_UP |
| 21 | LUCAS_HNF4A_TARGETS_UP |
| 22 | DAUER_STAT3_TARGETS_UP |
| 23 | DOANE_RESPONSE_TO_ANDROGEN_UP |
| 24 | KARLSSON_TGFB1_TARGETS_UP |
| 25 | ALONSO_METASTASIS_EMT_UP |
| 26 | PDGF_ERK_DN.V1_UP |
| 27 | HAN_JNK_SINGALING_UP |
| 28 | OZANNE_AP1_TARGETS_UP |
| 29 | ATF2_S_UP.V1_UP |
| 30 | RAFFEL_VEGFA_TARGETS_UP |
| 31 | MARKS_HDAC_TARGETS_UP |
| 32 | PRC2_EZH2_UP.V1_UP |
| 33 | HELLER_HDAC_TARGETS_UP |

**Table. S2.** Correlation values of SLC6A3 expression with signaling pathways regulation in ccRCC tumor.

| S. No. | Gene Signature | GSE40435 | GSE53757 |
|---|---|---|---|
| 1 | VANTVEER_BREAST_CANCER_ESR1_UP | 0.2373 | 0.2750 |
| 2 | WILLIAMS_ESR1_TARGETS_UP | -0.3414 | 0.0047 |
| 3 | YANG_BREAST_CANCER_ESR1_LASER_UP | 0.3488 | 0.2043 |
| 4 | KENNY_CTNNB1_TARGETS_UP | -0.1833 | -0.2342 |
| 5 | MANTOVANI_NFKB_TARGETS_UP | 0.1236 | 0.0137 |
| 6 | NGUYEN_NOTCH1_TARGETS_UP | -0.1152 | 0.3935 |
| 7 | KANNAN_TP53_TARGETS_UP | 0.0884 | -0.0651 |
| 8 | DER_IFN_ALPHA_RESPONSE_UP | -0.1490 | 0.0422 |
| 9 | DER_IFN_BETA_RESPONSE_UP | -0.1234 | 0.0694 |
| 10 | DER_IFN_GAMMA_RESPONSE_UP | -0.2191 | 0.0159 |
| 11 | AFFAR_YY1_TARGETS_UP | 0.1032 | 0.2368 |
| 12 | ELLWOOD_MYC_TARGETS_UP | -0.1709 | -0.0894 |
| 13 | LEE_LIVER_CANCER_MYC_UP | 0.2670 | 0.3583 |
| 14 | LEE_LIVER_CANCER_E2F1_UP | -0.0815 | -0.1377 |
| 15 | LANDIS_ERBB2_BREAST_TUMORS_65_UP | -0.2487 | -0.1593 |
| 16 | KRAS.LUNG_UP.V1_UP | 0.2020 | 0.2217 |
| 17 | IGARASHI_ATF4_TARGETS_UP | 0.1549 | 0.0229 |
| 18 | SRC_UP.V1_UP | 0.2377 | -0.3113 |
| 19 | ELVIDGE_HIF1A_TARGETS_UP | -0.3845 | -0.4240 |
| 20 | MCCLUNG_CREB1_TARGETS_UP | 0.2769 | 0.1282 |
| 21 | LUCAS_HNF4A_TARGETS_UP | 0.3449 | 0.3455 |
| 22 | DAUER_STAT3_TARGETS_UP | -0.1170 | -0.1345 |
| 23 | DOANE_RESPONSE_TO_ANDROGEN_UP | 0.1260 | -0.0672 |
| 24 | KARLSSON_TGFB1_TARGETS_UP | -0.3720 | -0.1794 |
| 25 | ALONSO_METASTASIS_EMT_UP | -0.1612 | -0.0156 |
| 26 | PDGF_ERK_DN.V1_UP | -0.1248 | -0.1361 |
| 27 | HAN_JNK_SINGALING_UP | 0.1570 | 0.0254 |
| 28 | OZANNE_AP1_TARGETS_UP | 0.0085 | 0.4650 |
| 29 | ATF2_S_UP.V1_UP | -0.2848 | -0.1654 |
| 30 | RAFFEL_VEGFA_TARGETS_UP | -0.1274 | -0.2009 |
| 31 | MARKS_HDAC_TARGETS_UP | 0.1048 | 0.2282 |
| 32 | PRC2_EZH2_UP.V1_UP | -0.1977 | 0.2687 |
| 33 | HELLER_HDAC_TARGETS_UP | 0.2606 | 0.0572 |

**Script File: S1**

Step 1:

```
d1<-read.csv("datasetname.csv", sep=",", header=TRUE)
genes<-d1[,1]
d2<-d1[,2:ncol(d1)]
d3<-scale(d2)
write.table(genes, file="output1_genes_input_file.txt")
write.table(d3, file="output2_scaled_input_values.txt", sep="\t")
```

Step 2:

```
d1<-read.csv("filename.csv", sep=",",header=TRUE)
sig<-read.csv("signaturefilename.csv", na.strings="NA", sep=",")

d1<-as.data.frame(d1)
sig<-as.data.frame(sig)
sig_head<-colnames(sig)

d1_colhead<-colnames(d1)

#write.table(d1_colhead, file="output_avg_signatures.txt", append=TRUE, sep="\t")

for(i in 1:length(sig_head)){
	signature<-sig[,i];
	signature<-as.matrix(signature);
	colnames(signature)="ID";
	siglic<-as.data.frame(signature)
	write.table(sig_head[i], file="output.txt",  append=TRUE, sep="\t")
	write.table(sig_head[i], file="output_avg_signatures.txt",  append=TRUE, sep="\t")
	d = merge(d1, siglic, by.x="ID", by.y="ID", incomparables=NA)

	d2<-d [,2:ncol(d)]
	Avg<-colMeans(d2)
	d_avg<-as.matrix(Avg)
	d_avg<-t(d_avg[,1])


#	stdev<-sd(d)
#	Avg<-mean(d)
	write.table(d, file="output.txt", append=TRUE, sep="\t")
```

```
	write.table(d_avg, file="output_avg_signatures.txt", append=TRUE, sep="\t")
}
```
